# Antimicrobial Resistance, Genomic, and Public Health Insights into Enterococcus spp. from Australian Cattle

**DOI:** 10.1101/2022.09.29.510230

**Authors:** Shafi Sahibzada, Rebecca Abraham, Terence Lee, David Jordan, Kate McMillan, Glen Mellor, Lesley Duffy, Mark O’Dea, Sam Abraham, Robert Barlow

**Affiliations:** Antimicrobial Resistance and Infectious Diseases Laboratory, Harry Butler Institute, Murdoch University, Western Australia, Australia; New South Wales Department of Primary Industries, Wollongbar, New South Wales, Australia; CSIRO Agriculture and Food, Coopers Plains, Queensland, Australia

## Abstract

Enterococci are opportunistic, potentially life-threatening pathogens of humans that are difficult to manage due to antimicrobial resistance. Historically, enterococci entering the food-chain through livestock have been viewed as a likely source of antimicrobial resistance in humans. Australian human-derived clinical enterococci have a propensity to be resistant to multiple classes of antimicrobials including vancomycin. Recent Australian studies involving pigs and chicken have virtually excluded these species as reservoirs of infection for resistant enterococci in humans. However, the Australian bovine population has not been similarly assessed. This study investigates the antimicrobial resistance profiles of enterococci from Australian cattle and the phylogenetic relationship between *E. faecium* isolated from cattle and human sepsis cases. Minimum inhibitory concentration assays were performed for *E. faecium* (n=343), *E. faecalis* (n=92), and *E. hirae* (n=284) against a panel of 15 antimicrobials. The majority of isolates were sensitive to all tested antimicrobials. Erythromycin resistance was most prevalent for *E. faecium* isolates (18.7%), daptomycin for *E. faecalis* (12.1%) and tetracycline for *E. hirae* (13.3%). Phenotypically, 1 *E. faecalis* was resistant to vancomycin and 9 were resistant to linezolid (*E. faecium* n=4, *E. faecalis* n=2, *E. hirae* n=3) but this was not confirmed with any know genotype. A subset of 67 *E. faecium* isolates selected for comparative phylogenetic analysis revealed that bovine isolates clustered with other livestock-derived and van-negative human isolates. In conclusion, there is a low prevalence of antimicrobial resistance amongst enterococci from Australian cattle which are unlikely to be precursor strains to vancomycin-resistant strains currently circulating in Australian hospitals.

**Importance:** *Enterococci* resistant to critically important antimicrobials such as vancomycin and linezolid are difficult to manage in health care settings. Historically, there has been the belief that livestock can act as a reservoir of resistance for human infections. Previous studies in poultry and pork have demonstrated that isolates derived from these livestock are unlikely to be precursor strains for currently circulating vancomycin resistant-*Enterococci* causing infection in humans. To date, there has been no study looking at the genetic similarity of bovine derived *Enterococci* and the relationship to human pathogenic strains. In this study we performed phenotypic and genotypic characterization of bovine derived *Enterococci* along with comparative phylogenetic analysis with other livestock derived and human sepsis derived *E. faecium* isolates. We found that while non-vancomycin resistant strain sharing occurs between hosts, cattle are unlikely to be precursor strains for vancomycin resistant human *E. faecium* infections.

## Introduction

Enterococci are commonly found as commensals of the gastrointestinal tract of humans and other animals but can behave as opportunistic pathogens, which are generally health care associated (Murray 1990, Lee, Pang et al. 2019). The majority of enterococci infections in humans are caused by *E. faecalis* and *E. faecium* resulting in mild urinary tract infections to severe sepsis (Lee, Pang et al. 2019). In the past three decades, there has been an increase in the prevalence of antimicrobial resistance (AMR) among enterococci, particularly among health care-associated enterococci infections resulting in adverse treatment outcomes (Carmeli, Eliopoulos et al. 2002). Globally, resistance to the critically important glycopeptide group, which includes vancomycin, has become a major public health issue (Lee, Pang et al. 2019, WHO 2019) and this is exemplified by high rates of vancomycin resistance found during surveillance of human enterococci in Australia (AURA 2021).

Resistance to vancomycin in enterococci is due to the presence of a *van* operon with *vanA* and *vanB* being the most clinically relevant (Werner, Coque et al. 2008, Freitas, Tedim et al. 2016). The *van* operon is of particular concern, as it is able to easily transfer between strains and even species, causing outbreaks to occur (Hegstad, Mikalsen et al. 2010). In America, the development of vancomycin resistance among enterococci is attributed to overuse of vancomycin in the health care setting (Leclercq 1997). However, in Europe it has been argued that vancomycin resistance first originated in livestock and became established in humans after zoonotic transfer although extensive studies of human outbreaks have failed to confirm this position (Aarestrup 1995, Bager, Madsen et al. 1997, Lester, Frimodt-Moller et al. 2006, Hammerum 2012).

In Australia it has been reported that 35.2% of *E. faecium* sepsis causing infections are vancomycin resistant. The *vanB* (60.6%) operon was more common than the *vanA* (38.8%) operon and a small proportion harboring both the *vanA* and *vanB* operons (0.6%) (Lee, Pang et al. 2020, Coombs, Daley et al. 2022). Surveillance of critical resistance in Australian food animals has not yet identified vancomycin resistance in enterococci due to *vanA* or *vanB* operons (Lee, Jordan et al. 2021). Although a study of cattle six years ago did not find any resistance to highly important antimicrobials (Barlow, McMillan et al. 2017) It is prudent to repeat this assessment to ensure emergence of critical resistance is detected promptly. This provides an opportunity for inaugural application of whole genome sequencing to enterococcus from Australian cattle to reinforce “one health” aspects of the ecology of enterococci in humans and animals.

In this study, a designed structured approach was used to assess the phenotypic and genotypic profile of AMR in enterococci isolated from cattle at slaughter from an ecological perspective. In addition, comprehensive genomic characterization was performed on a subset of *E. faecium* isolates from cattle. It was hypothesized that AMR prevalence would continue to be low, due to strict antimicrobial usage practices in Australia. We also hypothesized that cattle enterococci isolates would be genetically distinct from vancomycin resistant enterococci (VRE) that cause infection in humans.

## Methods

### Sample Collection

Faecal samples (40g) were collected 60 cm from the rectal end of the large intestine of cattle during the post-evisceration stage of processing. Samples were collected at random across four two-week sampling periods in February, March, June and August 2019. Proportionate stratified sampling based on the production class and slaughter volumes was applied to allocate the number of samples collected from each abattoir. A total sample collection target of 1000 was established, aiming to obtain 600, 200 and 200 samples from beef cattle, dairy cattle and veal calves, respectively.

Samples were stored chilled and returned to the laboratory via overnight courier. Samples were stored chilled for a maximum of 48 hrs before bacterial isolation. The work was approved by the Murdoch University animal ethics committee (Cadaver Permit No. 892).

#### Bacterial Isolation

Faecal samples were enriched using enterococcosel broth then streaked onto enterococcosel agar (BD, USA) and Slanetz and Bartley agar (Oxoid, UK). Enterococcosel agar plates were incubated at 35 ± 1°C for 24-48 h, whereas Slanetz and Bartley agar plates were initially incubated at 35 ± 1°C for 4h followed by 44 ± 1°C for 44 h. Following incubation, five presumptive *Enterococcus* colonies from each plate (10 colonies in total per sample) were patched onto SBA and incubated at 37 ± 1°C for 18 – 24 h. The resulting isolates were pooled into groups of five isolates and tested by PCR for the presence of *E. faecium* and *E. faecalis* using previously published protocols (Naserpour Farivar, Najafipour et al. 2014). If a pooled group of isolates tested positive for either *E. faecium* or *E. faecalis*, then further PCR testing of the individual isolates would occur. In samples that yielded both an *E. faecium* and *E. faecalis* isolate, both isolates were retained at -80°C using Protect bacterial preservers for antimicrobial susceptibility testing (AST). Isolates from samples that tested negative for *E. faecium* and *E. faecalis* were then confirmed as *Enterococcus spp*. by PCR (Dutka-Malen, Evers et al. 1995), identified to species level using MALDI-ToF and retained at -80°C using Protect bacterial preservers.

### Phenotypic antimicrobial susceptibility testing

Antimicrobial susceptibility testing (AST) was performed on all MALDI-ToF confirmed *E. faecium, E. faecalis* and *E. hirae* isolates by micro-broth dilution using custom made antimicrobial panels prepared on the RASP platform (Truswell, Abraham et al. 2021). Inocula were prepared manually and dispensed using an auto-inoculator. All phenotypic AMR assessments were completed following incubation at 37°C ± 1°C for 24 h. Isolates unable to display turbid growth in the growth control wells were considered invalid and repeated. Repeated isolates that failed to display turbid growth were removed from the analysis. *E. faecium* ATCC29212 was used as a control strain.

EUCAST ECOFF breakpoints were used to classify isolates into non-wild type (NWT) and wild types (WT) (EUCAST). NWT isolates (defined as having a minimum inhibitory concentration above the ECOFF) have been shown to contain acquired resistance mechanisms in their genome, even though they may have MICs below the defined CLSI clinical breakpoints. Therefore, for more simplistic determination of individual and multiclass resistance profiles, we refer to isolates exceeding the antimicrobial ECOFF as “resistant”. Multi-class resistant (MCR) isolates are therefore defined as having MICs above the ECOFF for one or more antimicrobial agents in three or more antimicrobial classes. This enables comparison with AMR surveillance systems such as DANMAP (https://www.danmap.org) and is more useful for assessing the recent emergence of resistance to multiple antimicrobial classes, particularly in populations expected to have low levels of resistance.

#### Genomic analysis

Whole genome sequencing was performed on all enterococci MCR isolates as well as isolates phenotypically resistant to daptomycin and vancomycin to determine presence of associated resistance genes. In Australia, *E. faecium* is of national importance as in the past up to 50% of human clinical isolates have been vancomycin resistant (VRE) (Australian Commission on Safety and Quality in Health Care 2019) and VRE are classed by WHO as a priority pathogen (Tacconelli, Carrara et al. 2018). Thus, a subset of 60 *E. faecium* isolates were selected for sequencing and genomic comparison. The isolates were chosen randomly by listing all un-sequenced *E. faecium* in the study and selecting every fifth isolate. DNA extraction was performed using the MagMAX Multi-sample DNA extraction kit as per the manufacturer’s instructions. DNA libraries were prepared using the Illumina Nextera XT DNA Library Prep Kit (Illumina, United States) according to the manufacturer’s protocol with an extended tagmentation time of seven minutes. Whole-genome sequencing (WGS) was performed on the Illumina NextSeq platforms using the NextSeq 500/550 Mid-Output Kit V2.5 (2×300 cycles). The raw sequence reads were de novo assembled using SPAdes V3.14.0 (Bankevich, Nurk et al. 2012). The genomes that passed the quality tests were used to predict the sequence type (ST) using the mlst tool V2.19 (Seemann 2020). Antimicrobial resistance genes and virulence genes were detected via ResFinder (Zankari, Hasman et al. 2012) and the universal virulence finder database VFDB (Liu, Zheng et al. 2019) using ABRicate V1.0.1 (Seemann 2021). The cut-off criterion for gene identity was 100% coverage and an identity over 95%. The serotyping prediction was performed on the WGS data using SeqSero2 V1.2.1 (Zhang, den Bakker et al. 2019).

Genome sequences were annotated with Prokka V1.14.6 (Seemann 2014, R Core Team 2021) and the pan-genome was extracted using Roary V3.13.0 (Page, Cummins et al. 2015). Principal component analysis (PCA) of binomial variables on the pan-genome matrix was performed for determination of association by total gene content and epigenetic traits. The 95% density ellipses were calculated (within R V4.0.5) (R Core Team 2021) from the specified correlation matrix (i.e., the first two components) and plotted with ggplot2 V3.3.3 (Wickham 2016). An approximate maximum likelihood phylogenetic tree was constructed for all isolates as on single nucleotide polymorphisms (SNPs) in the core genome utilizing the chromosomes of *Enterococci* obtained from NCBI. Core genes were aligned using SNIPPY V4.1.0 (Seemann 2015). Genomic data used for comparison was obtained from the AESOP from 2015-2017 (Lee, Pang et al. 2020) and previous studies in meat chickens (O’Dea, Sahibzada et al. 2019) and pigs (Lee, Jordan et al. 2021). All sequence data obtained from this study was deposited in the NCBI Sequence Read Archive under BioProject ID PRJNA858551.

### Statistical analysis

#### Phenotypic analysis

Confidence intervals of proportions were calculated using exact binomial confidence intervals using the Clopper-Pearson method. Significance of differences between enterprises in the proportion of isolates resistant to at least one antimicrobial (judged as P < 0.5) were assessed using Fisher’s Exact Test performed in Stata version 16.1 (StataCorp LLC, College Station, Texas USW, www.stata.com).

#### Genotypic analysis

Principal component analysis (PCA) of binomial variables on the pan-genome matrix was performed to determine the association between total gene content and epigenetic traits. The 95% density ellipses were calculated (within R) (R Core Team 2021) from the specified correlation matrix (i.e., the first two components) and plotted with ggplot2 V3.3.3 (Wickham 2016). Fisher exact tests were used to compare the proportion. Fisher exact methods were also used to calculate the confidence intervals for proportions.

## Results

### Sample collection

A total of 1001 faecal samples were collected across four sampling windows. Samples were collected from all three production classes; beef (n=591), dairy (n=194) and veal (n=216). Samples were collected from all Australian states with the majority of beef samples coming from Queensland (59.2%), the majority of dairy samples coming from Victoria (61.3%) and the majority of veal samples originating from NSW (47.2%) or Queensland (45.8%) establishments. Further analysis of samples sourced from beef cattle indicated that 285 (48.2%) were from grass-fed cattle, 235 (39.8%) were from feedlot cattle and 71 (12.0%) were from grain-assisted, grass-fed cattle.

### Enterococci isolation

*Enterococcus* was isolated from 546/591 (92.4%) beef cattle, 182/194 (93.8%) dairy cattle and 182/216 (84.3%) veal calf faecal samples for an overall detection of 90.9%. While the majority of samples yielded a single enterococcus isolate, 16 beef, 5 dairy and 3 veal samples yielded two isolates each, one *E. faecium* and one *E. faecalis*. A total of 9 different species of enterococcus were identified within the samples with *E. hirae* (42.5%) being the most prevalent species identified overall, followed by *E. faecium* (38.1%) then *E. faecalis* (9.9%). *E. mundtii* (4.3%), *E. casseliflavus* (4.2%), *E. gallinarum* (0.6%), *E. durans* (0.1%), *E. thailandicus* (0.1%) and *E. villorum* (0.1%) (TABLE 1).

**Table 1:** Prevalence of isolation of various *Enterococci* species from cattle (n=1001) at slaughter

| Species | Number of cattle (n=1001) | Percent (%) |
| --- | --- | --- |
| *E. hirae* | 398 | 42.5 |
| *E. faecium* | 357 | 38.1 |
| *E. faecalis* | 93 | 9.9 |
| *E. mundtii* | 40 | 4.3 |
| *E. casseliflavus* | 39 | 4.2 |
| *E. gallinarum* | 6 | 0.6 |
| *E. durans* | 1 | 0.1 |
| *E. thailandicus* | 1 | 0.1 |
| *E. villorum* | 1 | 0.1 |

### Phenotypic detection of AMR

All *E. faecium, E. faecalis* and *E. hirae* isolates were submitted for AMR analysis. A total of 719 isolates were able to grow under AMR test conditions and were included in the analysis, resulting in 343 *E. faecium*, 92 *E. faecalis*, and 284 *E. hirae* examined.

The majority of *E. faecium* isolates (72.8%) were sensitive to all antimicrobials tested (TABLE 2 and TABLE 5). Amongst *E. faecium* isolates no resistance to chloramphenicol, daptomycin, teicoplanin or vancomycin was observed. Erythromycin resistance (18.7%) was most prevalent. Resistance to gentamicin (0.6%), ampicillin (0.3%), penicillin (0.3%) and tetracycline (6.7%) was also observed. No ECOFF breakpoints are available for Quinupristin-Dalfopristin but 7.6% were clinically resistant based on CLSI breakpoints (Clinical and Laboratory Standards Institute 2021). Multi-class resistance (MCR – resistant to three or more antimicrobial classes) was observed in 1.5% of isolates with three isolates having a MCR profile of macrolides, streptogramins and tetracyclines, one isolate with a profile of beta-lactams, macrolides, streptogramins and tetracyclines and one isolate with a profile of oxazolidinones, macrolides, and tetracyclines. Beef cattle had the highest prevalence of resistance with 33.9% of isolates resistant to one or more antimicrobials. Variations in the prevalence of resistance to erythromycin across feed-type was observed with grain-assisted beef cattle having a prevalence of 56% resistance, feedlot beef cattle having a prevalence of 38.2% resistance and grass-fed cattle 8% resistance prevalence. Tetracycline and virginiamycin resistance was at least 2-3 times higher for feedlot cattle (17.6% and 23.5% respectively) when compared to grain assisted (8% for each drug) and grass fed (4.6% and 0% respectively). No ECOFF break-points are available for kanamycin, lincomycin and the concentration tested was above the ECOFF for streptomycin.

**Table 2:**
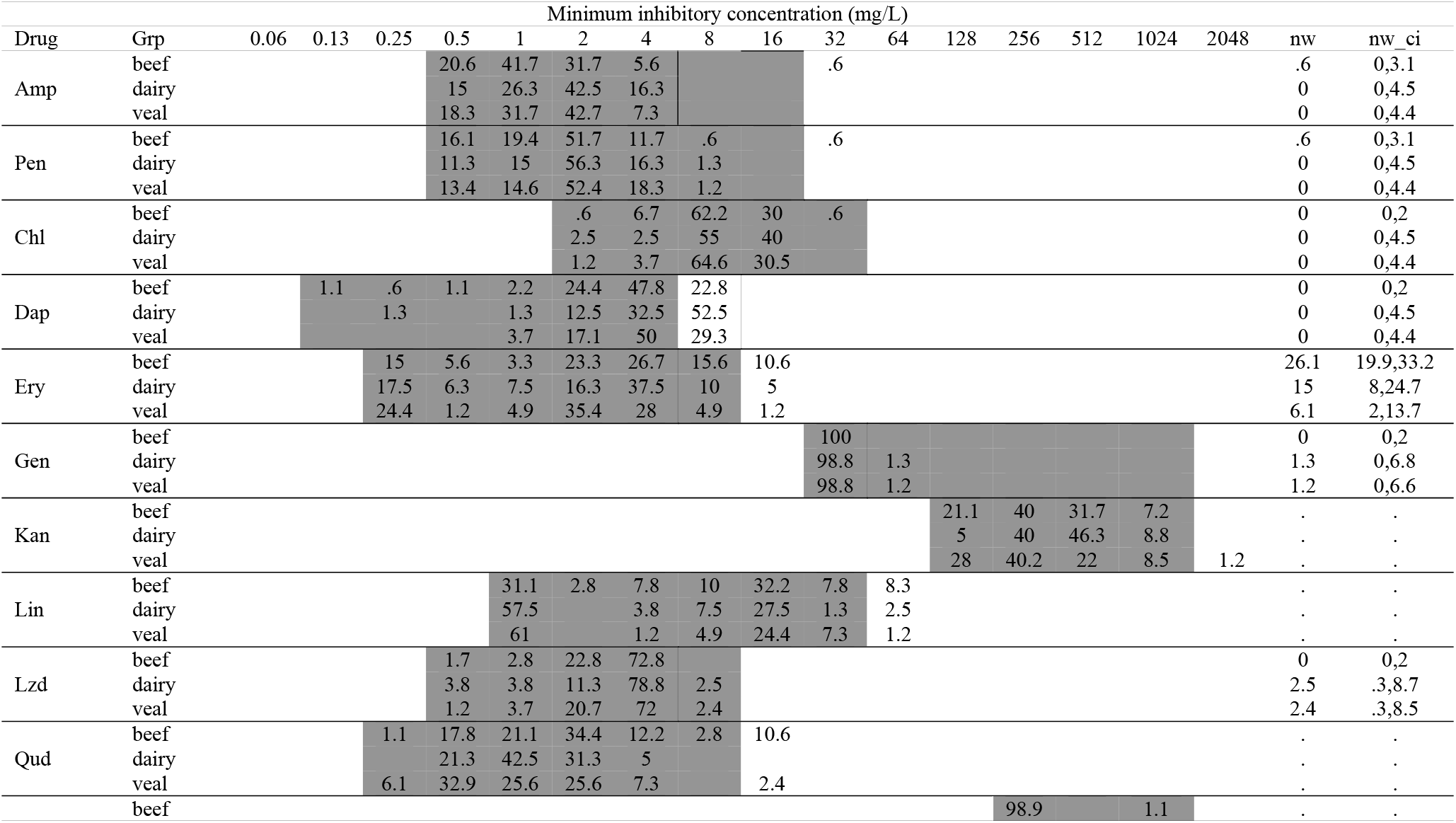
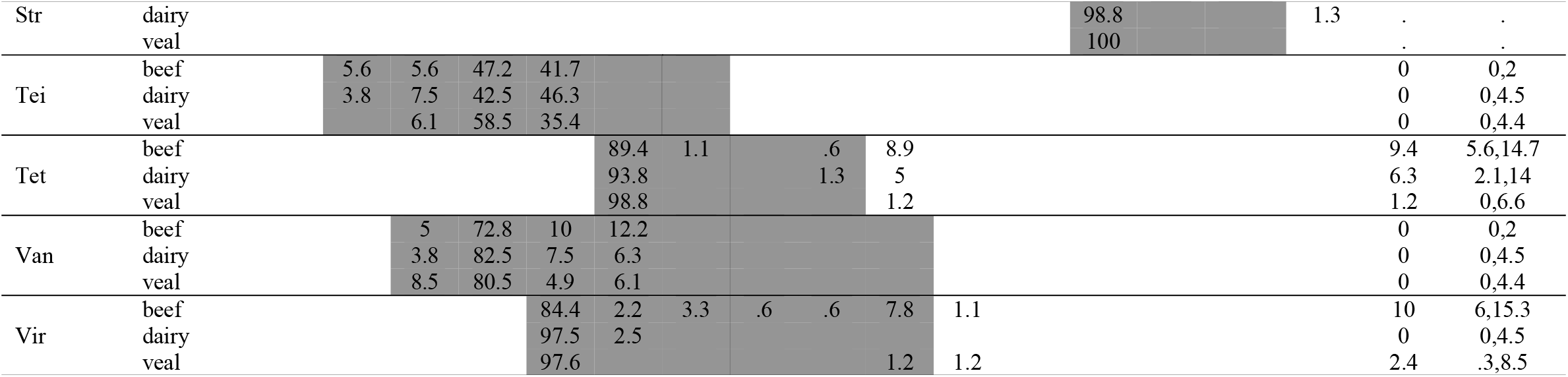
Distribution of minimum inhibitory concentration (MIC) for *E. faecium* isolated from Australian cattle assessed for sensitivity to 15 antimicrobials. Vertical bars represent the ECOFF non-wildtype breakpoints, shaded regions indicate the dilution range, numerical values are the percentage of isolates at each MIC. The percentage of isolates resistant (exceeding ECOFF) are presented under ‘nw’ with the corresponding 96% CI. The number of isolates for each animal group are beef (180), dairy (80) and veal (83). Amp = ampicillin, Pen = penicillin, Chl = chlorampehicol, Dap = daptomycin, Ery = erythromycin, Gen = gentamicin, Kan = kanamycin, Lin = lincomycin, Lzd = linezolid, Qud = quinupristin/dafopristin, Str = streptomycin, Tei = teicoplanin, Tet = tetracycline, Van = vancomycin, Vir = virginamycin

The majority of the *E. faecalis* isolates (79.1%) were susceptible to all antimicrobials tested (TABLE 3 and TABLE 5). There was no resistance to ampicillin, penicillin, gentamicin or virginiamycin detected. Low levels of resistance to chloramphenicol (1.1%), daptomycin (12.1%), erythromycin (1.1%), linezolid (2.2%), teicoplanin (1.1%), tetracycline (8.8%) and vancomycin (1.1%) was detected. Vancomycin and teicoplanin resistance was linked to a single MCR isolate which was also resistant to linezolid, isolated from a dairy cow. Only one other isolate had an MCR profile; macrolides, phenicols and tetracyclines, isolated from a veal calf.

**Table 3:**
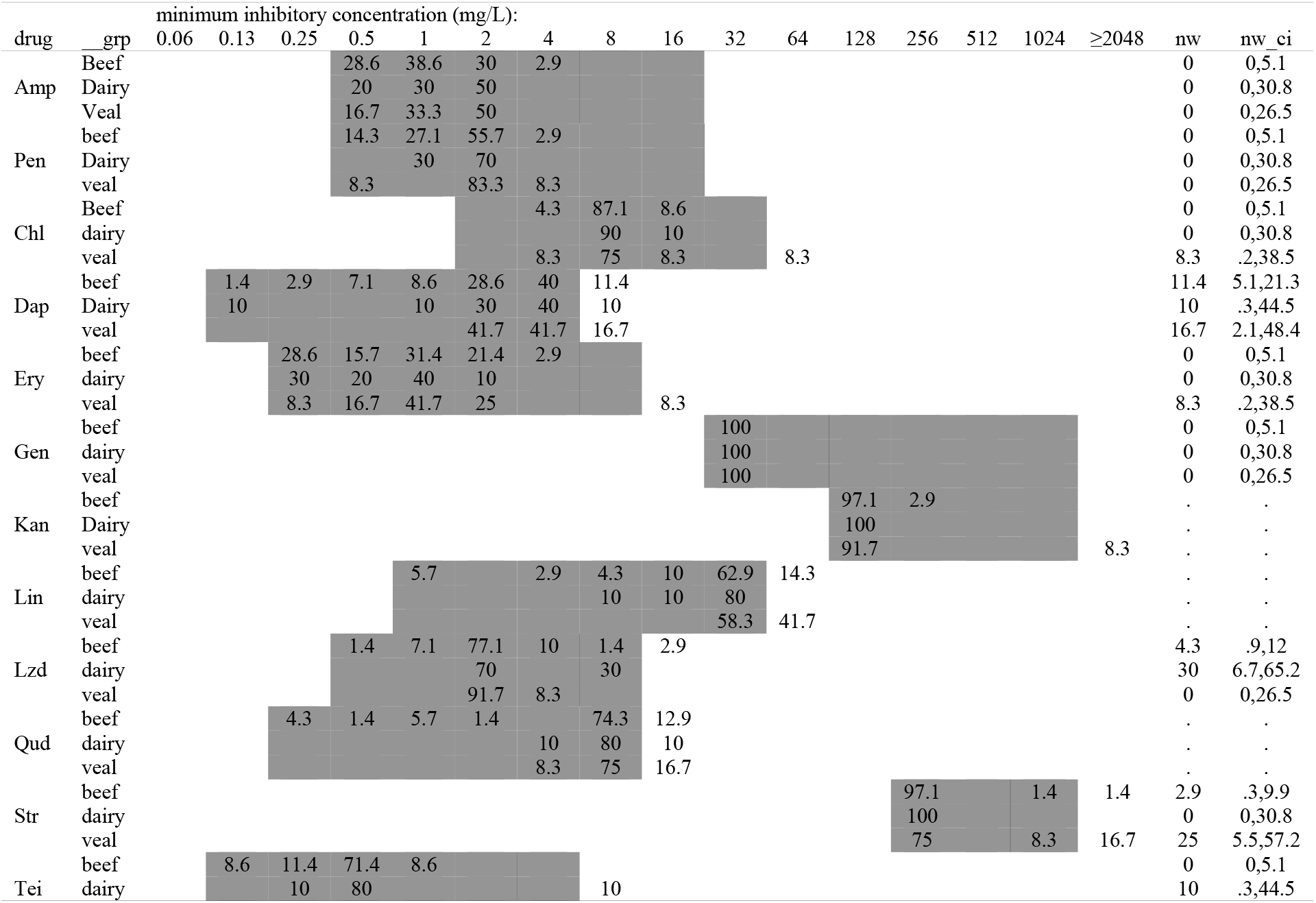
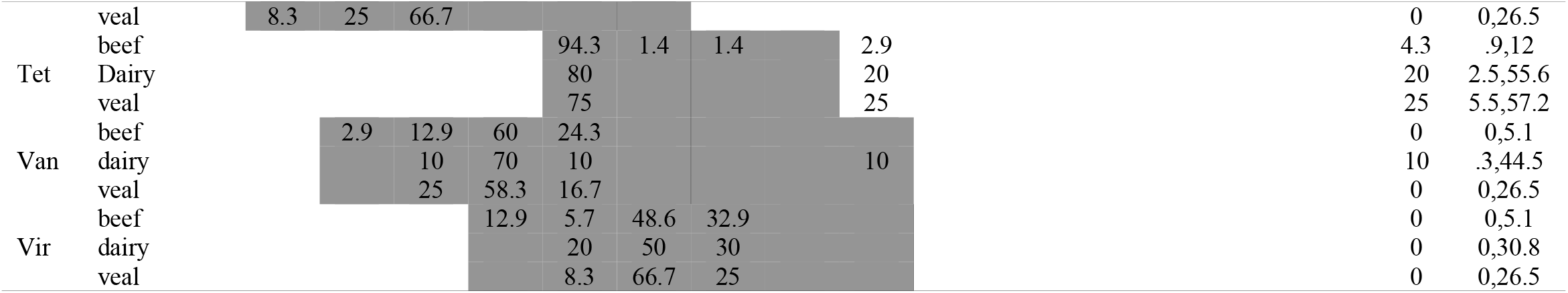
Distribution of minimum inhibitory concentration for *E. faecalis* isolated from Australian cattle assessed for sensitivity to 15 antimicrobials. Vertical bars represent the ECOFF non-wildtype breakpoints, shaded regions the dilution range, numerical values are the centage of isolates at each MIC. The percentage of isolates resistant (exceeding ECOFF) are presented under ‘nw’ with the corresponding 95% CI. The number of isolates for each industry are beef (70), dairy (10) and veal (12).

Although *E. hirae* was the most frequently isolated species in this study, only 71.9% of isolates could be tested for susceptibility due to a lack of growth under the test conditions. ECOFF values for *E. hirae* are only available for erythromycin, gentamicin and tetracycline. Resistance to erythromycin (6.6%), gentamicin (1.0%) and tetracycline (13.3%) was observed (TABLE 4). Using CLSI breakpoints clinical resistance to quinupristin-dalfopristin (5.6%) was also observed.

**Table 4:**
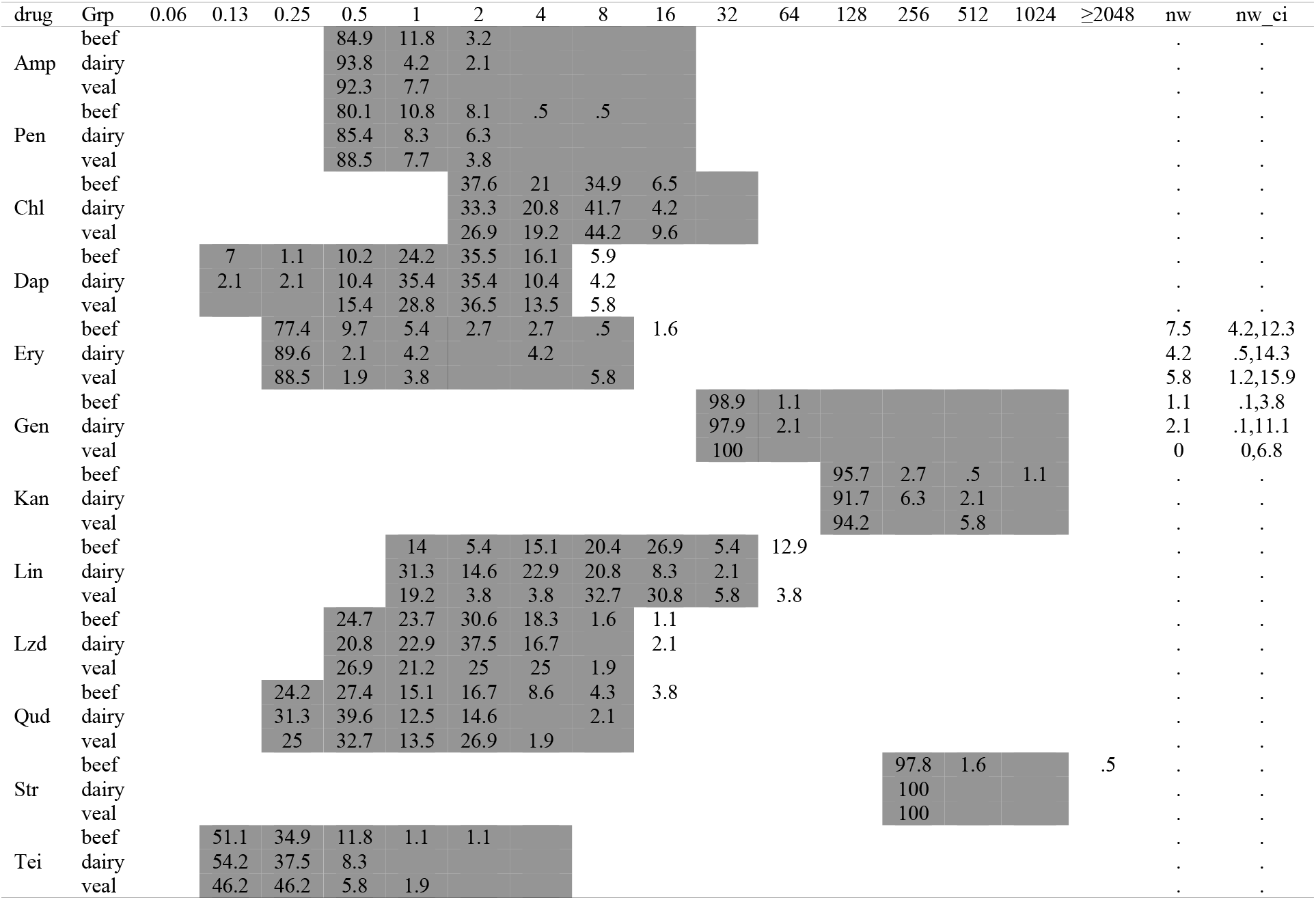
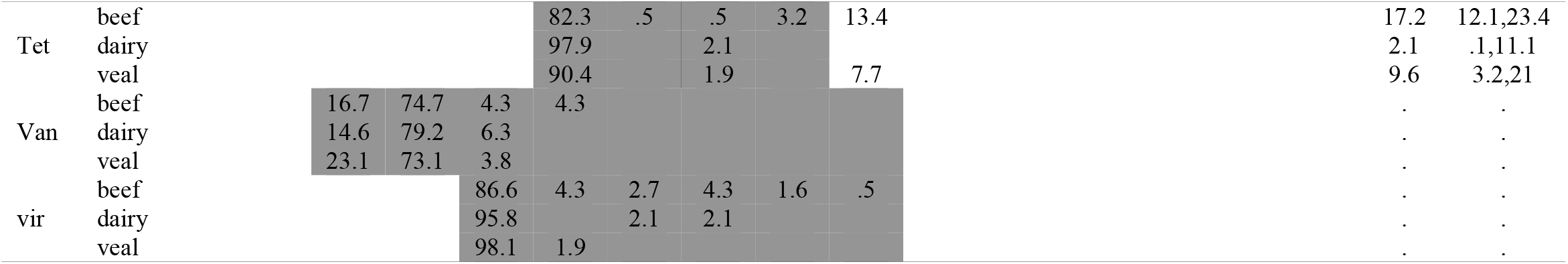
Distribution of minimum inhibitory concentration for *E. hirae* isolated from Australian cattle assessed for sensitivity to 15 antimicrobials. Vertical bars represent the ECOFF non-wildtype breakpoints, shaded regions the dilution range, numerical values are the percentage of isolates at each MIC. The percentage of isolates resistant (exceeding ECOFF) are presented under ‘nw’ with the corresponding 95% CI. The number of isolates for each industry are beef (185), dairy (48) and veal (51).

**Table 5:** resistance phenotype of *enterococci* isolated from Australian cattle.

|  | *E. hirae* |  |  | *E. faecium* |  |  | *E. faecalis* |  |  |
| --- | --- | --- | --- | --- | --- | --- | --- | --- | --- |
|  | Beef<br>(n=186) | Dairy<br>(n=48) | Veal<br>(n=52) | Beef<br>(n=180) | Dairy<br>(n=80) | Veal<br>(n=82) | Beef<br>(n=69) | Dairy<br>(n=10) | Veal<br>(n=12) |
| All sensitive | 131<br>(70.4)<sup>b</sup> | 42<br>(90.2) | 44<br>(84.6) | 116<br>(64.4) | 62<br>(77.5) | 71 (86.6) | 58<br>(84.0) | 7 (70) | 7<br>(58.3) |
| lip<sup>a</sup> | - | - | - | - | - |  | 8 (11.6) | - | 2<br>(16.7) |
| ami | 1 (0.5) | 1 (2.1) | - | - | 1 (1.3) | 1 (1.2) | - | - | - |
| mac | 7 (3.8) | 2 (4.2) | 3 (5.8) | 29 (16.1) | 11<br>(13.8) | 5 (6.1) | - | - | - |
| oxa | 1 (0.5) | 1 (2.1) | - | - | 1 (1.3) | 2 (2.4) | - | 1 (10) | - |
| str | 13 (7.0) | 1 (2.1) | - | 10 (5.6) | - | 2 (2.4) | - | - | - |
| tet | 24<br>(12.9) | 1 (2.1) | 5 (9.6) | 5 (2.7) | 4 (5.0) | 1 (1.2) | 3 (4.3) | - | 2<br>(16.7) |
| Lip tet | - | - | - | - | - | - | - | 1 (10) | - |
| Ami tet | 1 (0.5) | - | - | - | - | - | - | - | - |
| Mac str | 1 (0.5) | - | - | 8 (4.4) | - | - | - | - | - |
| Mac tet | 5 (2.7) | - | - | 6 (3.3) | - | - | - | - | - |
| Oxa tet | 1 (0.5) | - | - | - | - | - | - | - | - |
| Str tet | - | - | - | 2 (1.1) | - | - | - | - | - |
| Gly oxa tet | - | - | - | - | - | - | - | 1 (10) | - |
| Mac oxa tet | - | - | - | - | 1 (1.3) | - | - | - | - |
| Mac phe tet | - | - | - | - | - | - | - | - | 1 (8.3) |
| Mac str tet | 1 (0.5) | - | - | 3 (1.7) | - | - | - | - | - |
| Bla mac str tet | - | - | - | 1 (0.6) | - | - | - | - | - |

<sup>a</sup> lip – lipopeptides, ami – aminoglycosides, mac – macrolides, oxa – oxazolidinones, str – streptogramins, tet – tetracyclines, gly – glycopeptides, phe – phenicols and bla – beta-lactams. <sup>b</sup> figures in parenthesis are percent. – no profile present for species/industry.

### Genotypic analysis of resistance

MCR *E. faecium* (n=5) and *E. faecalis* (n=2) isolates as well as a random subset of *E. faecium* (n=60), were included for genomic analysis to identify MLST, virulence, resistance genes and pan genome. Both *E. faecalis* sequenced were MCR phenotypes. The first isolate was ST147 with genotypic resistance genes *dfrE* and *Isa(A)*. It was phenotypically resistant to vancomycin, linezolid and tetracycline but no known genes conferring these resistances were identified. The second isolate, with a phenotypic resistance profile of macrolides, phenicols and tetracyclines was ST623 with genotypic and phenotypic resistance matching.

MLST analysis of 65 *E. faecium* isolates (5 MCR and 60 subset) identified 24 different sequence types amongst 34 isolates and 31 isolates with unknown STs. The most common MLST for *E. faecium* was ST32 (n=7) followed by ST214 (n=3) and ST22 (n=3). The remaining STs present had a single isolate identified.

The *msrC* gene that belongs to the efflux pump gene family and confers resistance to erythromycin and other macrolide and streptogramin B antibiotics was found in 76.1% (n=51) of *E. faecium* isolates. Another macrolide associated resistance gene, *erm(B)*, was also identified in 6.0% (n=4) of *E. faecium* isolates. The pleuromutilin resistance gene *eat(A)* was identified in 76.1% (n=51) of the isolates. Aminoglycoside resistance genes, *aac*(6’)-Ii (94.2%, n=63) and *spw* (1.52%, n=1), were also noted. Tetracycline resistance genes, *tet*(L), *tet*(M), and *tet*(S), were found in 3-9.1% of isolates. A *vat(E)* genotype was found in a small number of isolates (3.0%, n=2).

### Comparative genomic analysis

Comparative genomic analysis of 1218 *E. faecium* isolates was performed. All sequenced *E. faecium* (n=65) isolates from this study were compared to 1024 isolates from human sepsis cases in Australia collected between 2017-2019 and routine surveillance isolates from pig fecal material (n=69) (Lee, Jordan et al. 2021) or chicken caecal material (n=60) (O’Dea, Sahibzada et al. 2019). A SNP based maximum likelihood phylogenetic tree showed that most human isolates with *van* genes present cluster together. All cattle isolates cluster separately with chicken and pig isolates, and some of the human isolates that are negative for the *van* operon (FIGURE 1).

**Figure 1:**
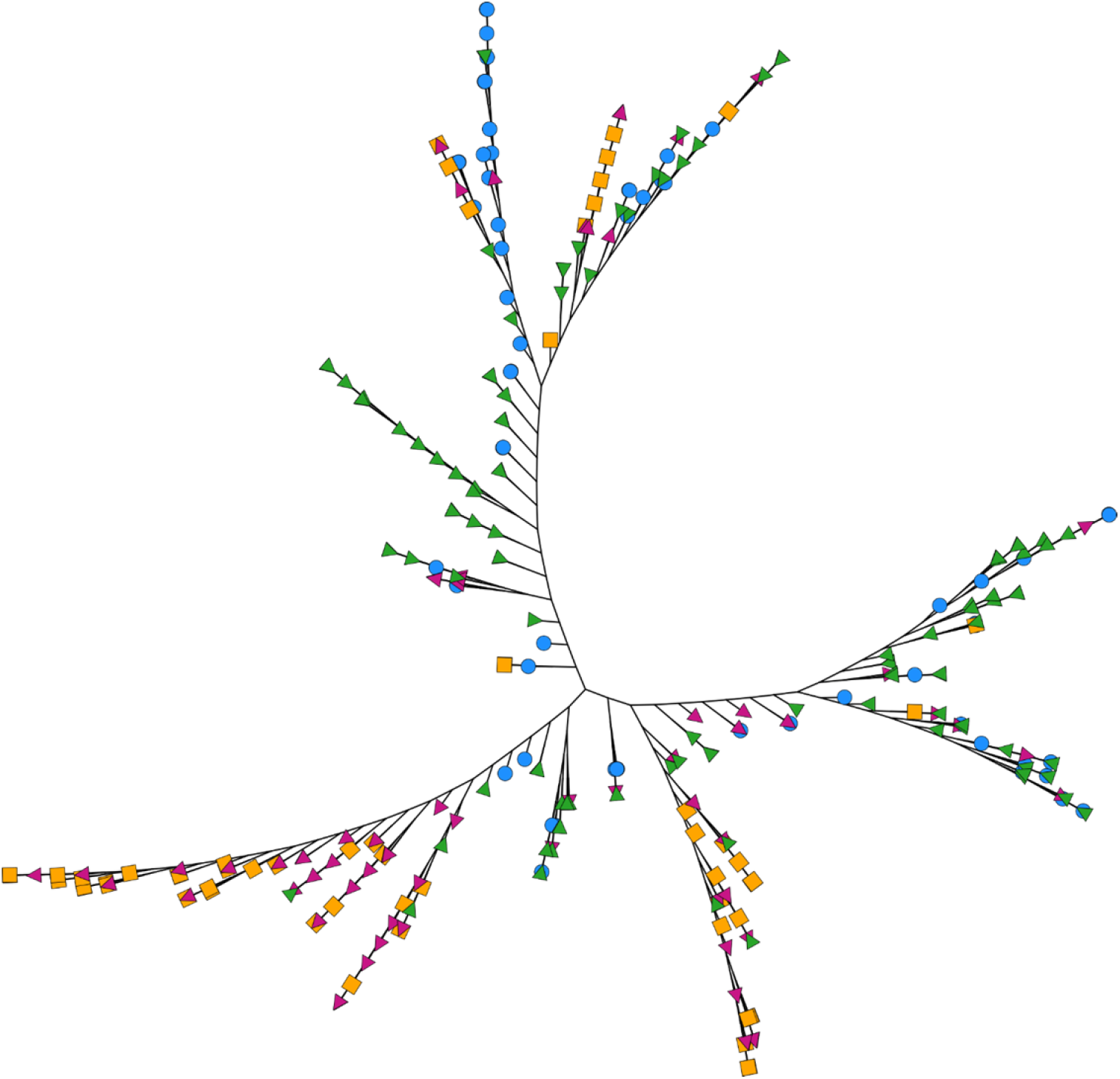
Maximum-likelihood phylogenetic tree based on single nucleotide polymorphisms. The tree includes *E. faecium* isolates from cattle (n=65, blue circles), pigs (n=69, pink right-pointing triangle), chickens (n=60, orange square) and human sepsis isolates that clustered with the animal isolates (n=115, green left-pointing triangle). Branches are not to scale. Figure was annotated using iTOL.

Principal component analysis was also performed on total gene content with 44,043 genes identified. Based on the presence or absence of the vancomycin-resistant genes (*vanA, vanB*), the 95% density ellipses identified three clusters comprised of *vanA* positive, *vanB* positive, and *vanA/B* negative. The *vanA* positive and *vanB* positive isolates were mutually exclusive based on 95% density ellipse. All cattle isolates were clustered with *van*-negative isolates from animals and humans (FIGURE 2).

**Figure 2:**
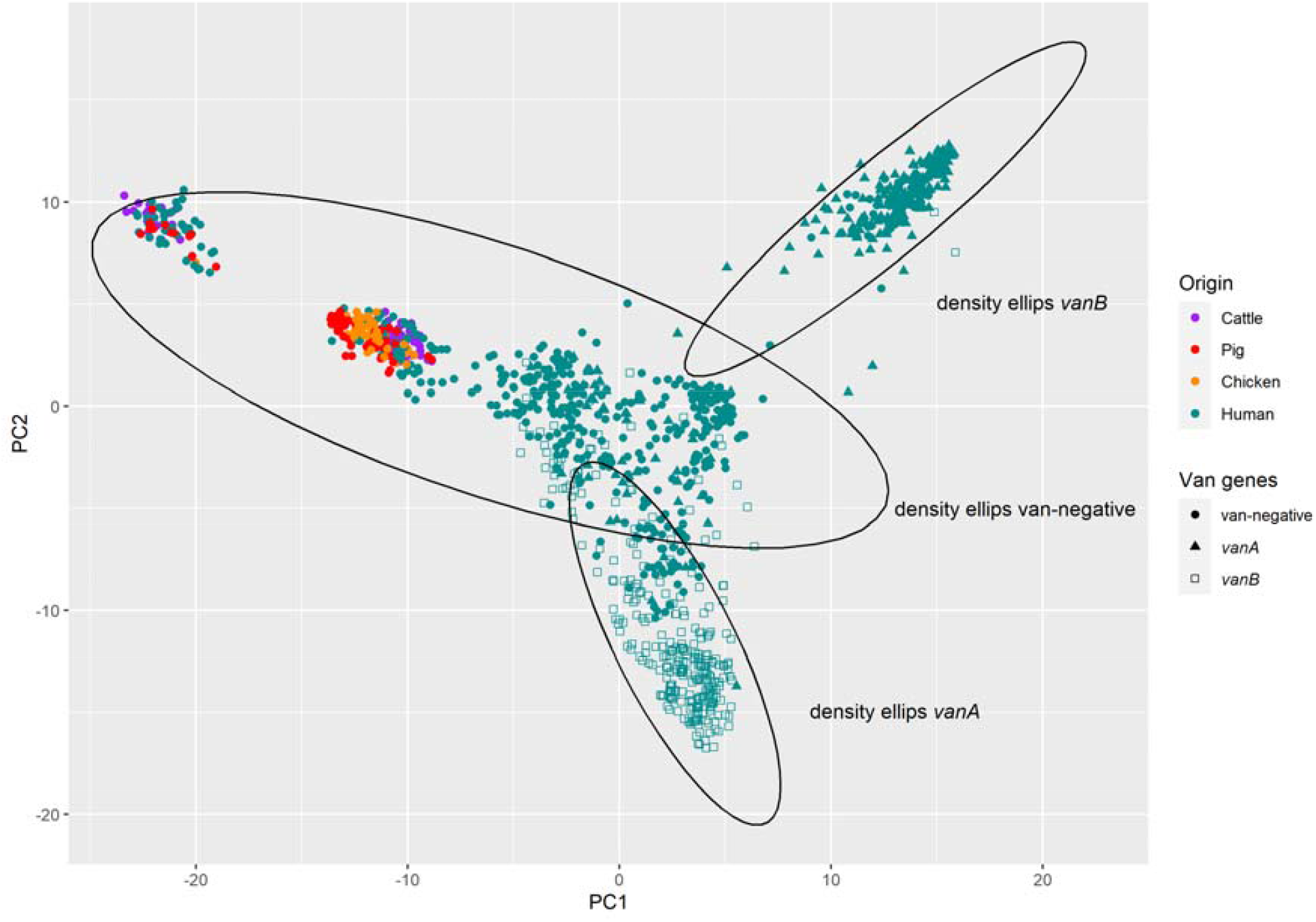
A principal component analysis of total gene content of *E. faecium* isolates from cattle (n=65), pigs (n=69), chicken (n=65) and human sepsis (n=1024). Isolates harboring the *van*A operon (solid triangles), *van*B operon (hollow squares) and *van*-negative (solid circles) are drawn with a 95% density ellipse.

## Discussion

In this study we investigated the occurrence and phenotypic characteristics of enterococci isolated from cattle, focusing on the three most dominant species identified, *E. hirae, E. faecium* and *E. faecalis*. We continued with genotypic characterization focused on the two most important species, *E. faecium* and *E. faecalis* (Lee, Pang et al. 2019) as these are standard components of national surveillance systems and have implications for public health arising from zoonotic transfer Data from this study on phenotypic and genotypic analysis of bovine enterococci supports the hypothesis that resistance to antimicrobials, in particular CIA, is low. The majority of isolates in this study were sensitive to all antimicrobials tested (74.7%). Globally, resistance in *Enterococci* from cattle varies considerably. In similar studies from The United States of America, South Korea and countries within the European Union, including Denmark, France and Italy, rates of clinical resistance in *E. faecium* and *E. faecalis* from cattle origin to critically important antimicrobials vancomycin and linezolid was 0-0.3%. Prevalence of resistance to erythromycin amongst *E. faecium* was 3.6%, 19.9% and 9.0%; tetracycline 14.2%, 25.9% and 5.2%; quinopristin/dalfopristin 65.6%, 8.7% and 2.2%; daptomycin 2.8%, 5.6% and 0% respectively. Prevalence of clinical resistance in *E. faecium* from Australian cattle was low; erythromycin 7%, tetracycline 6.1%, quinopristin/dalfopristin 7.6% and daptomycin 0% (de Jong, Simjee et al. 2019, Food and Drug Administration 2021, Kim, Moon et al. 2021). Internationally prevalence of resistance in *E. faecalis* to erythromycin between 2.4 -20.4%, tetracycline 20 – 41.8% and daptomycin 0 – 1.3%. In this study prevalence of clinical resistance was low; 1.1% for erythromycin, 7.7% for tetracycline and 0% for daptomycin (de Jong, Simjee et al. 2019, Food and Drug Administration 2021, Kim, Moon et al. 2021).

Phenotypic resistance to CIAs vancomycin, teicoplanin and linezolid were initially each identified however phenotypic results were not supported by genotypic analysis with no known glycopeptide or oxazolidinone resistance genes being identified in phenotypically resistant isolates. The elevated MIC of all linezolid resistant *E. faecalis* and *E. faecium* are likely false-positives due to the phenomenon of MIC drift, as the MICs are only one dilution higher than the ECOFF break-point (Abraham, O’Dea et al. 2019). The *optrA* gene, reportedly conferring resistance to linezolid and phenicols, was identified in a single *E. faecalis* isolate which was phenotypically resistant to chloramphenicol but sensitive to linezolid (Wang, Lv et al. 2015).

An MCR profile was observed in a small number of cases (1.1%). Feed-type did not seem to affect presence of MCR in beef cattle. A higher prevalence of virginiamycin resistant *E. faecium* isolates amongst feedlot beef cattle isolates (23.5%, n=16/68) compared to grass-fed beef isolates (0%, n=0/87) could be due to the use of virginamycin to prevent ruminal acidosis during grain feeding (Meat & Livestock Australia 2019). Resistance to both streptogramins tested, virginiamycin (ECOFF break points) and quinupristin/dalfopristin (CLSI break points), was identified in the majority of streptogramin resistant isolates (89.7%) while quinupristin/dalfopristin resistance alone was observed in the remaining isolates. Of the streptogramin resistant *E. faecium* isolates sequenced (n=8), one had the *eat(A)* gene mutation only, five had the *msr(C)* and *eat(A)* genes and one had the *msr(C), eat(A)* and *erm(B)* genes which confer resistance to streptogramin B and macrolides, while the final isolate had all three previously mentioned genes in addition to a *vat(E)* gene which confers resistance to streptogramins A (Miller, Munita et al. 2014). Overall, the results are consistent with the previous Australian study in 2013 with the exception of macrolide and streptogramin resistance which have both increased in *E. faecium* since 2013 (Meat & Livestock Australia 2019).

A genomic comparison focused on *E. faecium* identified four STs associated with human and porcine *van-*negative sepsis isolates; ST22, ST32, ST94 and ST116 which are not typical with hospital strains (Lee, Jordan et al. 2021). Although isolates from cattle are unlikely to be the source of CIA resistant infections in humans, sharing of STs between isolates from cattle, humans and pigs suggests evidence of bi-directional transmission, coevolution or a common exposure source between these three species (Lee, Jordan et al. 2021). Principal component analysis (PCA) of the total gene content further supported the genetic similarity between cattle, pig, chicken and van-negative human isolates (FIGURE 2). The majority of livestock isolates clustered together in the PCA, with a smaller cluster consisting of human, cattle and porcine isolates only. However, the maximum likelihood tree based on core genes indicated that the majority of cattle isolates were more similar to human derived van-negative isolates, clustering more frequently with human derived isolates. Cattle derived isolates were not found dispersed with the majority of porcine and poultry derived isolates which formed two main clusters (FIGURE 1). This analysis supports the hypothesis that human clinical isolates in Australia represent a distinct subset not closely related to cattle or other livestock derived isolates.

Antimicrobial resistance in enterococci isolated from Australian cattle was low and sequenced isolates clustered only with van-negative human and other livestock derived isolates, however, there are several limitations to this study. Firstly, we were only able to study 1-2 representative enterococci per sample. This sampling strategy does not capture the phenotypic or genotypic diversity of enterococci within a sample, potentially missing profiles of low prevalence. Due to the sampling strategy, we are also unable to comment on the distribution of strains and species within each sample. Secondly, genomic characterization of the isolates from other sources were from previously available data from historic Australian collection from humans (diagnostic cases or routine infection control screenings) and other animal species from AMR surveys and this may pose a sampling and interpretation biases. However, the current study provides insight into the relationship between *E. faecium* isolated from cattle and clinical isolates from humans, suggesting the majority of isolates are not closely related.

In conclusion, this study has provided significant insights into the characteristics of three commonly identified enterococci species from Australian cattle. No genetic resistance to the CIAs of human medicine vancomycin, teicoplanin and linezolid were identified. Comparison of the genomics of *E. faecium* isolated from cattle was not genetically similar to human clinical isolates baring *van* resistance operons. This study supported previous Australian studies in various livestock species indicating that food-producing animals are not a reservoir for vancomycin resistant *E. faecium* or vancomycin resistant *Enterococci* genetic elements in Australia.

---

**Supplementary Figure 1:**
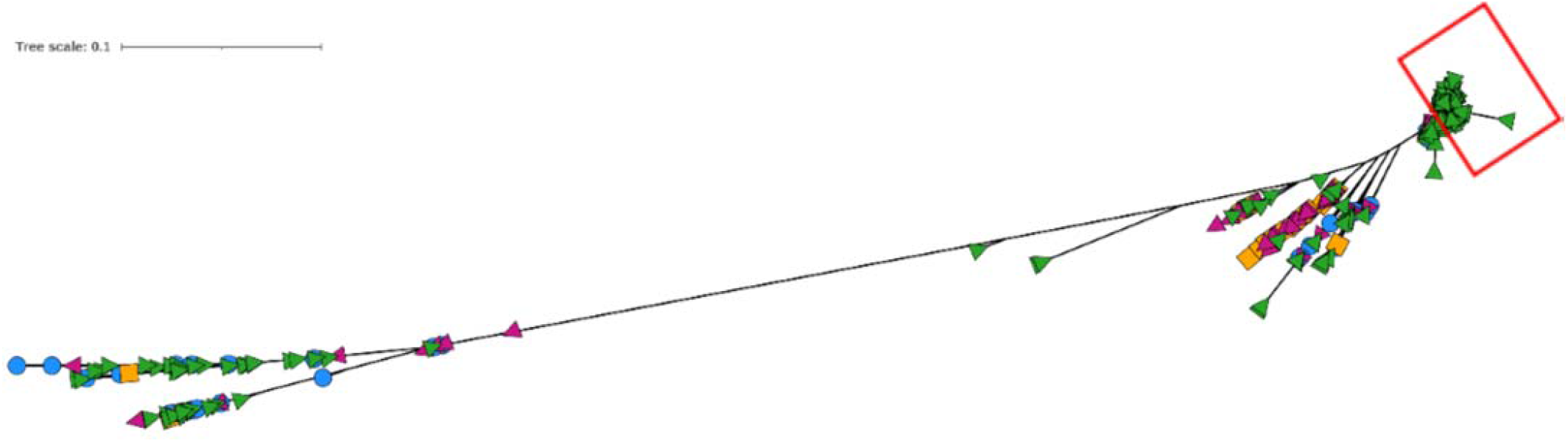
maximum-likelihood phylogenetic tree based on single nucleotide polymorphisms. The tree includes 1218 *E. faecium* isolates from cattle (n=65, blue circle), pigs (n=69, pink right-pointing triangle), chickens (n=60, orange square) and human sepsis isolates (n=1024, green left-pointing triangle). The phylogenetic tree was constructed using 709 core-SNPs found across 1054 core genes. All strains from the current study along with other food animal isolates from Australia are clustered within the *van* negative and separate from the *van* positive (red box) (*van*A and *van*B).

